# Untargeted Metabolomics to Expand the Chemical Space of the Marine Diatom *Skeletonema marinoi*

**DOI:** 10.1101/2023.09.07.556696

**Authors:** Mahnoor Zulfiqar, Daniel Stettin, Saskia Schmidt, Vera Nikitashina, Georg Pohnert, Christoph Steinbeck, Kristian Peters, Maria Sorokina

## Abstract

Diatoms (Bacillariophyceae) are aquatic photosynthetic microalgae with an ecological role as primary producers in the aquatic food web. They account substantially for global carbon, nitrogen, and silicon cycling. Elucidating the chemical space of diatoms is crucial to understanding their physiology and ecology. To expand the known chemical space of a cosmopolitan marine diatom, *Skeletonema marinoi*, we performed High-Resolution Liquid Chromatography-Tandem Mass Spectrometry (LC-MS^2^) for untargeted metabolomics data acquisition. The spectral data from LC-MS^2^ was used as input for the Metabolome Annotation Workflow (MAW) to obtain putative annotations for all measured features. A suspect list of metabolites previously identified in the *Skeletonema* spp. was generated to verify the results. These known metabolites were then added to the putative candidate list from LC-MS^2^ data to represent an expanded catalogue of 1970 metabolites estimated to be produced by *S. marinoi*. The most prevalent chemical superclasses, based on the ChemONT ontology in this expanded dataset, were “Organic acids and derivatives”, “Organoheterocyclic compounds”, “Lipids and lipid-like molecules”, and “Organic oxygen compounds”. The metabolic profile from this study can aid the bioprospecting of marine microalgae for medicine, biofuel production, agriculture, and environmental conservation. The proposed analysis can be applicable for assessing the chemical space of other microalgae, which can also provide molecular insights into the interaction between marine organisms and their role in the functioning of ecosystems.

**Importance:** Diatoms are abundant marine phytoplankton members and have great ecological importance and biochemical potential. The cosmopolitan diatom *Skeletonema marinoi* has become an ecological and environmental research model organism. In this study, we used untargeted metabolomics to acquire a general metabolic profile of *S. marinoi* to assess its chemical diversity and expand the known metabolites produced by this diatom. *S. marinoi* produces a chemically diverse set of secondary metabolites with potential therapeutic properties, such as anti-cancer, antioxidant, and anti-inflammatory. Such metabolites are highly significant due to their potential role in drug discovery and bioeconomy. Lipids from *S. marinoi* also have potential in the biofuel industry. Furthermore, the environmental fluctuations in the water bodies directly affect the production of different secondary metabolites from diatoms, which can be key indicators of climate change.

## 1 Introduction

Unicellular microalgae, forming the phytoplankton, are primary producers in the marine food web, contributing to biodiversity and affecting the biogeochemical cycles, particularly the oceanic carbon cycling [1, 2]. Among the phytoplankton community, diatoms (Bacillariophyceae) are found in the euphotic layer of the ocean and have the largest contribution to the aquatic biosphere [3]. These eukaryotic microalgae survive under extreme environmental conditions such as varying light, temperature, nutrition, and salinity. They produce unique Natural Products (NP) with distinct biological activities displaying various chemical classes [4]. As the primary producers, diatoms serve as a substantial biological source that can generate diverse natural products with applications in various NP-driven industries, such as food technology, cosmetics, biofuel, nanotechnology, and drug discovery [5–8]. However, the marine diatom community remains a large reservoir of unknown natural products [9].

To expand the record of identified marine natural products or metabolites, we used *Skeletonema marinoi* (NCBI Taxonomy ID: 267567) as a model organism for diatoms. The strain used was taxonomically revised from *Skeletonema costatum* [10]. *S. marinoi* is a marine unicellular, centric diatom that lives in chain-like colonies consisting of up to 30 cells, with 1-2 chloroplasts in each cell located near the siliceous cell wall or frustule [10].

*Skeletonema marinoi* is considered a non-toxic species and generally safe [11]. *S. marinoi* has been evolutionary successful, and its genome encodes certain bacterial, plant, and animal-like metabolic pathways [12]. It can also adapt to varying environmental conditions [13]. Given the optimal availability of nutrients, *S. marinoi* produces massive phytoplankton algal blooms in the coastal regions [14], releasing oxylipins that act as toxins for predators and other phytoplankton members [15, 16]. Although much effort has been made for the chemical characterisation of lipids in *S. marinoi*, as they impact growth, reproduction, stress response, and defense mechanism in the diatoms [17–20], much of its metabolome is still unexplored compared to other model species.

Mass spectrometry (MS) based metabolomics is the key technology for detecting these secondary metabolites and covers the analysis of compounds with distinct structural heterogeneity [21]. Here, we performed the metabolome annotation of the mass spectrometry data obtained from *S. marinoi* to comprehend the chemical composition of the primary and specialised metabolites produced by the diatom. We annotated molecular structures, chemical classes, and molecular formulae to MS features obtained with High-resolution Liquid Chromatography-Electrospray Ionization-Tandem Mass Spectrometry (LC-ESI-Orbitrap MS^2^) using Metabolome Annotation Workflow (MAW) [22]. The findings of this study contribute to the expansion of known marine natural products derived from *S. marinoi*.

## 2 Results

### 2.1 Suspect List of Known *S. marinoi* Metabolites

A suspect list of known metabolites produced by *S. marinoi* and the closely related *S. costatum*, was created using different databases and literature articles, described in the methodology section. The curated suspect list contains names, formulae, species, SMILES, InChIs, monoisotopic masses, PubChem identifiers, source databases and references, IUPAC names, synonyms, subclasses, classes, and superclasses of 893 compounds. For the classification of these metabolites, ClassyFire [23] was used, which annotated 851 compounds from the suspect list. ClassyFire uses ChemONT, which is a structure-based taxonomy system used to annotate chemical compounds to a hierarchical system of classification. Metabolites belong to the class “Organic compounds” and can be categorised into more detailed superclasses (generic), classes (specific), and subclasses (highly specific). The most abundant superclasses in the suspect list, according to ChemONT, were “Organoheterocyclic compounds”, “Organic acids and derivatives”, “Organic oxygen compounds”, and “Lipids and lipid-like molecules” (see Figure S1 from supplementary materials).

To comprehend the structural similarity among the known metabolites from the suspect list, we clustered together the chemically similar compounds from the abundant superclasses into a network (shown in Figure S2 in supplementary materials). This chemical similarity network visually represents the structural similarity between different compounds, which can reveal joint functional regulation. A group of compounds in a cluster can represent a biochemical pathway or otherwise connected regulations. The chemical similarity network was created using an “all vs all” approach, where all chemical structures in the suspect list were compared against one another using a Tanimoto similarity score threshold of >= 0.85. The chemical superclass with the highest internal similarity in the suspect list was the Lipids and lipid-like molecules, exhibiting the polyunsaturated fatty acids (PUFA), polyunsaturated aldehydes (PUA), and precursor molecules such as acetyl CoA and malonyl CoA which drive the citric acid cycle and fatty acid biosynthesis [24]. Organoheterocyclic compounds were also highly interconnected representing different tetrapyrroles such as chlorophyll a. The other two superclasses were amino acid derivatives and coenzymes, and organic oxygen compounds belonging to the polyphenols.

### 2.2 Structure Annotation and Classification of Tandem Mass Spectrometry Data from *S. marinoi*

To broaden the known chemical space of *S. marinoi*, we acquired untargeted metabolomics data from *S. marinoi* samples using High-Resolution Liquid Chromatography-Tandem Mass Spectrometry (HR-LC-MS^2^). The data was measured in both positive and negative modes, which resulted in 1014 and 839 features, respectively, making a total of 1853 distinct metabolic features with unique precursor masses [*m*/*z*] and retention time (seconds), out of which 1153 were identified as unique metabolites. This difference in unique features and unique compounds can be attributed to the presence of various isotopes, adducts, and fragments from the same compound [25]. The Metabolome Annotation Workflow (MAW) was used to perform dereplication using spectral databases (GNPS [26], Massbank [27], and HMDB [28, 29]), and compound databases integrated with SIRIUS4 [30, 31]. The results were post-processed and each distinct feature was assigned a list of putative structures with Metabolomics Standards Initiative (MSI) confidence levels of identification [22, 32, 33]. Table 1 shows the number of unique features annotated within each MSI level.

**Table 1:** Number of features from the LC-MS^2^ data of *S. marinoi*, annotated with MAW, divided into different MSI levels.

| MSI level | Description | No. of features |
| --- | --- | --- |
| 1 | Structure identification with a standard | No standards were used here, the results for standards with MAW can be found in [22] |
| 2 | Experimental spectral matching with one or more spectral databases | 147 |
| 3 | Tentative candidate structure | 1130 |
| 4 | Molecular formula or chemical class | 4 |
| 5 | Only the spectral features from LC-MS<sup>2</sup> | 572 |

To assess the metabolic profile of *S. marinoi* from LC-MS^2^ data, the high-scoring candidates with MSI levels 2 and 3 were analysed. The annotations contained both primary metabolites such as amino acids, and their derivatives (pipecolic acid, ornithine, carnitine), nucleic acids and analogs (adenosine, guanosine, cyclic AMP), carbohydrates (alpha-lactose and glucose-1-phosphate); and fatty acids (eicosapentaenoic acid, myristic acid, stearic acid), and different other organic acids (gluconic acid, chlorogenic acid, and ascorbic acid). The composition of compound superclasses, classes, and subclasses in this experimental dataset is represented by a sunburst plot in Figure S3 in supplementary materials. A total of 1036 annotations were assigned a chemical superclass using CANOPUS [34] and ClassyFire. This dataset shows the same abundant superclasses as the suspect list.

### 2.3 Expansion of known metabolome of *S. marinoi*

Lastly, the putative candidate list from the LC-MS^2^ data was compared with the suspect list to extract metabolites that were common in both datasets with tanimoto similarity score of 1, which led to 57 common compounds. The most prevalent class was carboxylic acids and derivatives, particularly amino acids and analogues, as most of the common metabolites were primary metabolites (Figure S4 in supplementary materials). The compounds from the suspect list and LC-MS^2^ data were then combined in a single union set of metabolites, which resulted in 1997 compounds. Manual curation was performed to remove expectedly misannotated metabolites originating from plant, fungal, or animal origins which then reduced the list to 1970 compounds that represent the expanded chemical space of compounds produced by *S. marinoi*. Figure 1 represents a sunburst plot of the chemical classification of 1691 compounds from this expanded chemical space of compounds from *S. marinoi*.

**Figure 1:**
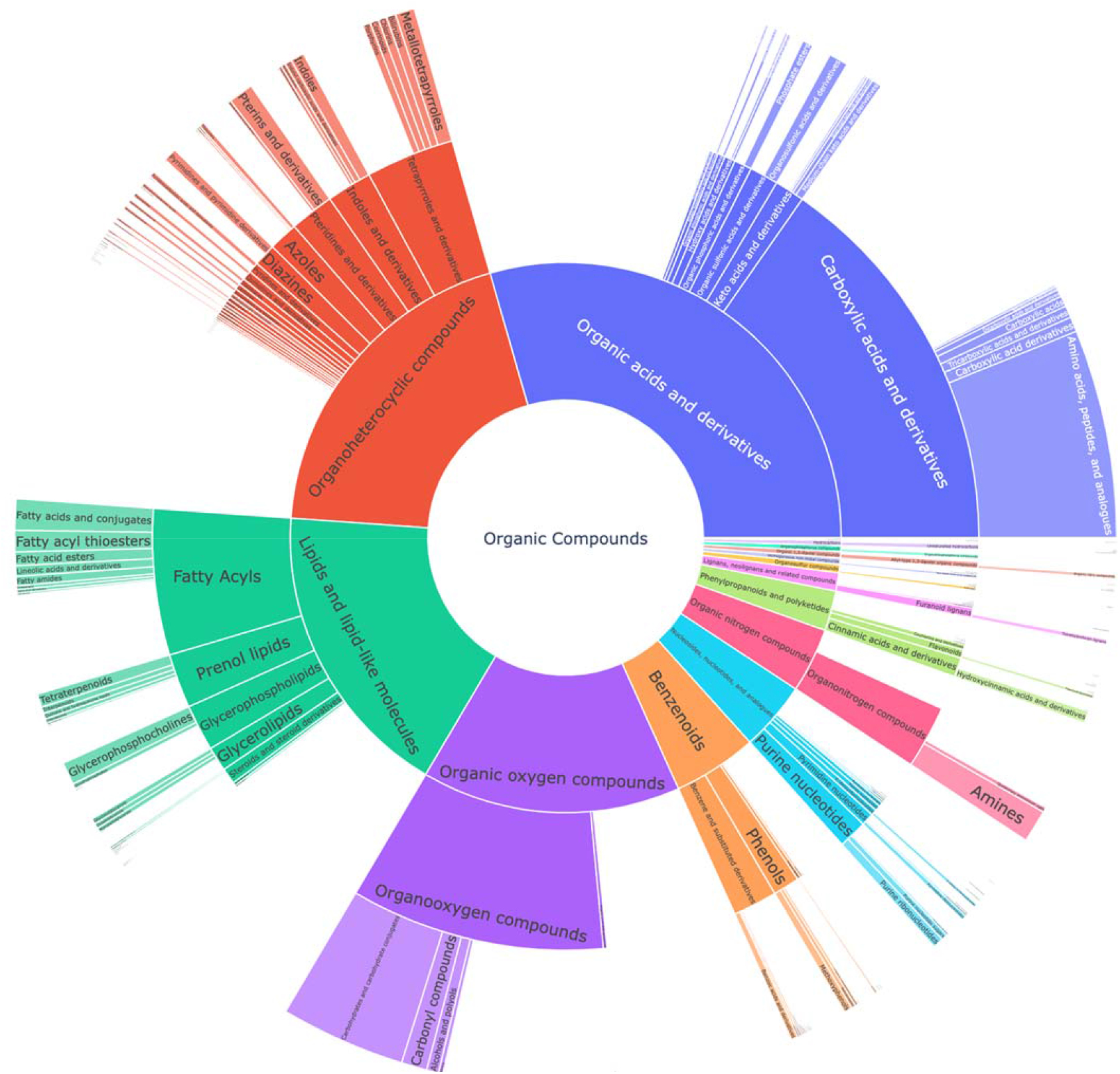
Sunburst plot for chemical classification of the union set from the suspect list and the LC-MS^2^ data from *S. marinoi* which shows the expanded known metabolome of the diatom. The innermost section of the kingdom of organic compounds is classified into superclasses, classes, and subclasses. The most prevalent superclasses are Organic acids and derivatives (28.1%), Organoheterocyclic compounds (20.3%), Lipids and lipid-like molecules (17.5%), and Organic oxygen compounds (14.5%).

The metabolites from the LC-MS^2^ data were incorporated into the chemical similarity network of compounds from the suspect list, shown in Figure S2 in supplementary materials, to demonstrate the expansion in the chemical similarity, which indicated the metabolites previously not reported or missing from the chemical similarity network or the known metabolic pathways of the diatom (as depicted in Figure 2). The hexagonal-shaped metabolite nodes in Figure 2 represent the putative candidates from the LC-MS^2^ data. Figure 2 shows that the “Lipids and lipid-like molecules” superclass had again the largest expansion representing newly added metabolites such as saturated oxo-fatty acid (SOFAs), fatty acid amides (FAAs), and glycerophospholipids. The “Organic oxygen compounds” superclass expanded with building blocks of polysaccharides for the diatom cell wall. Cyanodecanoic and cyanododecanoic acids were added to the chemical similarity network of “Organic acids and derivatives” superclass.

**Figure 2:**
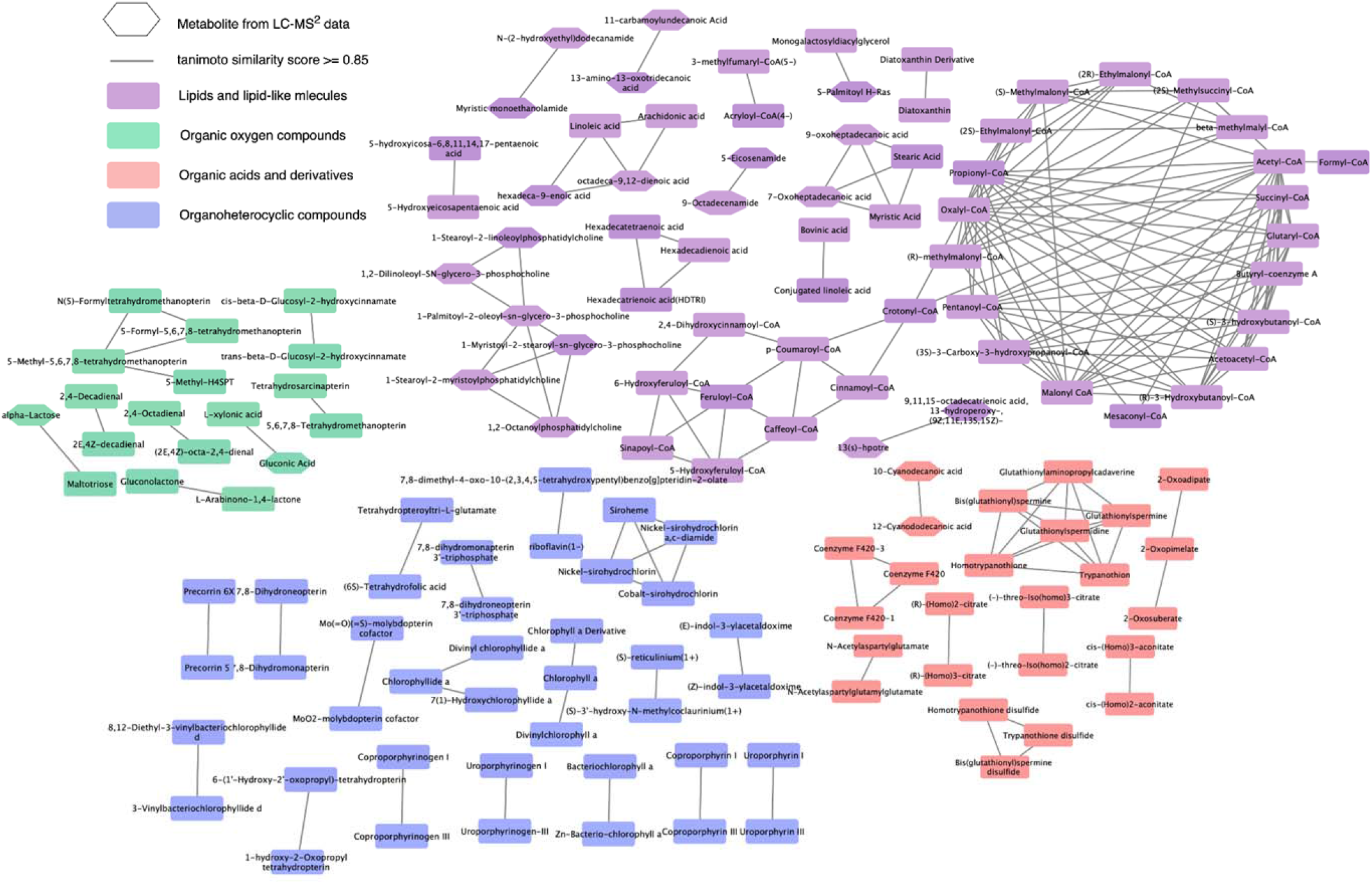
Expansion of chemical similarity network of the suspect list with the candidate structures from LC-MS^2^ data. This network represents the expanded chemical similarity space of the suspect list from Figure S2 in the supplementary materials; the new metabolites from the LC-MS^2^ data are represented with a hexagonal-shape This network shows the most abundant superclasses and the structural similarities between the metabolites from these superclasses.

## 3 Discussion

Marine natural product reservoirs are still partly untapped [9]. The marine diatom *S. marinoi*, for example, has been extensively studied for its lipid metabolism, however, the chemical diversity has not yet been fully explored for other natural products. To understand the biochemical mechanisms occurring within ubiquitous diatoms, we expanded the chemical space of known metabolites from *S. marinoi*, using a whole untargeted metabolomics approach. With our automated Metabolome Annotation Workflow (MAW), we annotated 69% of the metabolic features from the LC-MS^2^ data, with a chemical structure or a molecular formula (see Table 1). We verified the presence of previously identified metabolites using the suspect list of compounds from *S. marinoi* as the source of known metabolites (see Supplementary Figures S1 and S2). Here we discuss metabolites, annotated in this study, of which, some have not been previously identified in *S. marinoi*.

### 3.1 Secondary metabolites in the expanded chemical space of *S. marinoi*

Using the Liquid Chromatography-Tandem Mass Spectrometry (LC-MS^2^) data, MAW annotated some of the known metabolites produced by *S. marinoi*, many of which were amino acids, analogues, and their derivatives such as betaine, pipecolic acid, 4-hydroxyproline, methionine sulfoxide, and N,N-dimethyl arginine. Such polar compounds alleviate under salt stress in *S. marinoi* and might act as osmoprotectants [35]. *S. marinoi* is also known to produce a rich portfolio of “Lipids and lipid-like molecules”. With MAW, we annotated several polyunsaturated fatty acids (PUFA) like eicosapentaenoic acid and eicosatrienoic acid, saturated fatty acids (SFA) such as myristic acid and stearic acid, and fatty acid esters (FAE) such as isovalerylcarnitine, propionylcarnitine and butyrylcarnitine, which also act as stress protectants [35]. Being a photosynthetic organism, *S. marinoi* is also known to produce terpenoid pigments, such as astaxanthin, and tocopherol, which were also verified with MAW [11].

MAW annotated 1072 unique structures to metabolites that were not present in the suspect list and hence represented the expansion of known metabolites produced by *S. marinoi*. Only a few of these compounds show chemical similarity with the previously known natural products produced by *S. marinoi* (shown in Figure 2 as hexagonal nodes), among which most of the metabolites belonged to the “Lipids and lipid-like molecules” superclass. Lipids in diatoms have been studied extensively due to their role as membrane constituents, regulators of gene expression, and sources of energy for the cells via fatty acid beta-oxidation [36, 37]. In Figure 2, one of the chemical similarity clusters showed two saturated fatty acids (SFA), myristic (14:0), and stearic acids (18:0), and two saturated oxo-fatty acids (SOFA) 7-oxoheptadecanoic acid and 9-oxoheptadecanoic acid (derivatives of heptadecanoic acid). SOFAs are SFA with a ketone group. This remains an understudied class of lipids with unknown pathways [38] and hence cannot be linked back to the known diatom metabolism. However, the SFA heptadecanoic acid is present in algae, plants, and bacteria [39]. In a study done on *S. costatum*, the fatty acid profile in the stationary growth phase showed minute levels of heptadecanoic acid [19] indicating the measurable presence of this SFA after the end of the cell growth period. Heptadecanoic acid has been exploited for its medicinal use against type 2 diabetes and skin cancer [40, 41]. 4-Oxo sebacic acid which is likely derived from sebacic acid (a dicarboxylic SFA) was also annotated using MAW, however, no previous records of sebacic acid in diatoms have been reported so far.

Another class of detected lipids is the fatty acid amide (FAA). A chemical similarity pair was formed between two FAA derivatives of myristic and lauric acid: Myristic monoethanolamide and N-(2-hydroxyethyl)dodecanamide respectively. Both are N-functionalised long-chain-acyl ethanolamines resulting from the condensation of the carboxy group of an SFA with the amino group of ethanolamine. N-(long-chain-acyl)ethanolamines are converted to SFAs by FAA hydrolases [42]. Another pair of FAAs with high structural similarity was 5-eicosenamide (an unsaturated eicosanoic acid derivative) and 9-octadecanamide (stearic acid derivative). Two other high-scoring examples (from Table 2) are hexadecanamide (derived from palmitic acid) and tetradecanamide (derived from myristic acid). Carboxamides also undergo conversion back to SFA. SFAs are present in the lipid profile of many diatoms [43], however, there is no systematic research done on their FAA derivatives reported in this study. FAAs, such as oleamide from oleic acid, have anti-biofouling effects [44] on the accumulated microalgal biofilms on various surfaces, but this result has to be verified in the light of our finding that microalgae contribute themselves to the compound class.

**Table 2:**
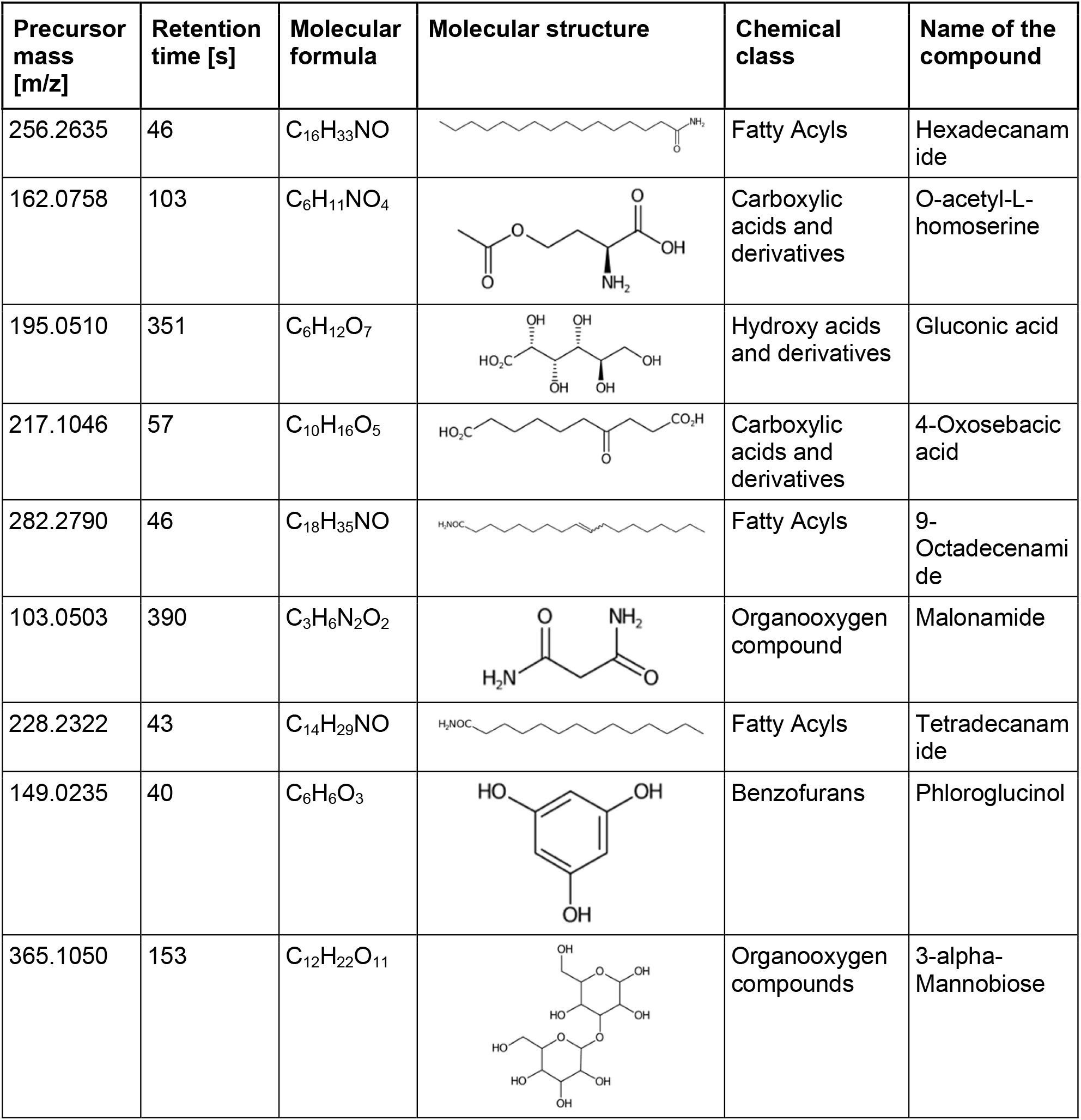
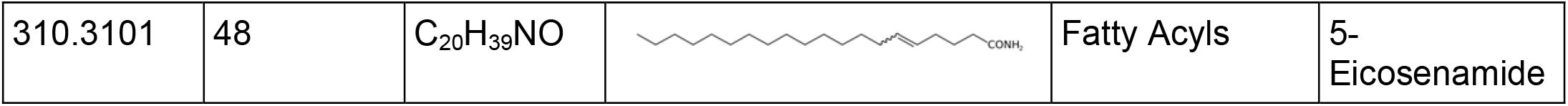
List of top-ranking metabolites (based on MAW candidate selection criterion) from the LC-MS^2^ data annotated with MAW from MSI Levels 2 and 3. **Figure 3:** Workflow for untargeted mass spectrometry approach to acquire the whole metabolome from *Skeletonema marinoi*. After the sample collection and metabolic extraction, LC-MS^2^ measurements are performed using an orbitrap mass spectrometer. The data obtained from the LC-MS^2^ is then subjected to MAW which performs the annotation of metabolites and generates a final list of metabolites extracted from the samples.

| Precursor mass [m/z] | Retention time [s] | Molecular formula | Molecular structure | Chemical class | Name of the compound |
| --- | --- | --- | --- | --- | --- |
| 256.2635 | 46 | C<sub>16</sub>H<sub>33</sub>NO |  | Fatty Acyls | Hexadecanamide |
| 162.0758 | 103 | C<sub>6</sub>H<sub>11</sub>NO<sub>4</sub> |  | Carboxylic acids and derivatives | O-acetyl-L-homoserine |
| 195.0510 | 351 | C<sub>6</sub>H<sub>12</sub>O<sub>7</sub> |  | Hydroxy acids and derivatives | Gluconic acid |
| 217.1046 | 57 | C<sub>10</sub>H<sub>16</sub>O<sub>5</sub> |  | Carboxylic acids and derivatives | 4-Oxosebacic acid |
| 282.2790 | 46 | C<sub>18</sub>H<sub>35</sub>NO |  | Fatty Acyls | 9-Octadecenamide |
| 103.0503 | 390 | C<sub>3</sub>H<sub>6</sub>N<sub>2</sub>O<sub>2</sub> |  | Organooxygen compound | Malonamide |
| 228.2322 | 43 | C<sub>14</sub>H<sub>29</sub>NO |  | Fatty Acyls | Tetradecanamide |
| 149.0235 | 40 | C<sub>6</sub>H<sub>6</sub>O<sub>3</sub> |  | Benzofurans | Phloroglucinol |
| 365.1050 | 153 | C<sub>12</sub>H<sub>22</sub>O<sub>11</sub> |  | Organooxygen compounds | 3-alpha-Mannobiose |
| 310.3101 | 48 | C<sub>20</sub>H<sub>39</sub>NO |  | Fatty Acyls | 5-Eicosenamide |

One larger cluster in the chemical similarity network in Figure 2 is the group of six glycerophospholipids: mainly phosphocholines and phosphatidylcholines with two fatty acyl groups. Glycerophospholipids are integral to the membrane structure [45] and have also been found in the silica deposition vesicles that form the silica shells in the diatom *Thalassiosira pseudonana* [46]. Enzymatic transformation of these metabolites releasing choline, occurs through hydrolysis via phospholipase D, where choline acts as a downstream signaling molecule [47]. Another reaction is the acyltransferase reaction which catalyses the release of fatty acids from the 1 or 2 acyl group position via phospholipase A [48]. These enzymatic activities have also been reported in diatoms [49, 50], but not with the glycerophospholipids found in this work. It is noteworthy that the spectral features annotated with these glycerophospholipids had smaller precursor masses as compared to the monoisotopic mass of the molecules which can be attributed to the in-source fragmentation where one fatty acyl chain fragmented from the whole molecule, and so the annotations in this case are not extremely reliable.

Diatoms produce extracellular polysaccharides as part of their silica shell/ frustules formed by monosaccharide building blocks. Generally, the cell wall of diatoms contains also mannose and glucuronic acid as main constituents and varying amounts of fucose and xylose [51]. A recent study showed that the diatom *Phaeodactylum tricornutum* cell wall contains α -(1 → 3)-mannan backbone which could explain the presence of 3-alpha-mannobiose in diatoms as annotated with MAW [62]. Furthermore, xylose, maltotriose, and allulose were annotated with MAW and had a chemical similarity with three sugar molecules from the suspect list, arabinose, maltose, and sorbose, respectively with a tanimoto similarity score of 1. MAW annotated maltotriose which is found in pennate diatom biofilms [52]. Another monosaccharide allulose was annotated with MAW which is rarely found in natural carbohydrate plants and macroalgae, with applications such as anti-atherosclerosis, anti-hyperlipidemic, and neuroprotective [53].

A few other metabolites annotated with MAW were: phloroglucinol, and malonamide, and two polyphenols (chlorogenate and 4-hydroxycinnamic acid). Phloroglucinol is a growth promoter produced by marine algae *Phaeophyceae* and *Fucaceae*, and is known to increase the levels of the carotenoid pigment fucoxanthin in microalgae [54]. Malonamide is a dicarboxylic acid diamide, a derivative of malonic acid. Malonamide can be converted back to malonic acid via amidase. Malonamide derivatives (MAMDs) have been utilised as anticancer and antibiotic drugs [55], however, not many studies have been conducted on the role of malonamide as a metabolite specifically in diatoms.

Diatoms in general are rich sources of polyphenolic compounds exhibiting antioxidant properties [56]. Studies show that diatom *Phaeodactylum tricornutum* under copper stress, releases high concentrations of polyphenols such as 4-hydroxycinnamic acid and chlorogenic acid [57, 58].

### 3.2 Prokaryote-associated metabolites

*S. marinoi* forms synergistic interactions with marine prokaryotes. The phycosphere of the marine diatom, which represents the diffusion limited mucus containing layer around the cells, hosts several bacterial species which stimulate the growth of the diatoms [59]. The bacteria provide the *S. marinoi* with vitamins and organic sulfur metabolites and get the nitrogen and carbon sources in return [60, 61]. Since, in this study, *S. marinoi* cultures (non-axenic) were not isolated from their indigenous prokaryotic associates, bacterial-produced metabolites were also measured and annotated. Here, we discuss a few high-scoring metabolites annotated with MAW reported in marine prokaryotes.

A sugar-based organic acid, gluconic acid has been found in marine bacteria connected to the release of domoic acid as a toxin in blooms of the diatom Pseudo-nitzschia multiseries [62]. A benzenoid from the statin family which is known for its therapeutic properties, fluostatin J, is generally found in marine actinobacteria [63]. Indole is produced by symbiotic bacteria supporting the growth of Skeletonema marinoi [59]. Marine bacteria and some marine algae execute the biodegradation of sulfoacetate to produce sulfoacetaldehyde, which is a major organic solute in marine organisms [64]. 6-Chloropurine riboside derived from the bacterium Geobacillus stearothermophilus, is used as an antiviral and anti-tumoral therapeutic agent [65].

### 3.3 Assessment of data quality and metabolome annotation workflows

The metabolites annotated with MAW revealed a large number of chemical compounds from *Skeletonema marinoi*, which have been also previously reported in other diatoms. However, we also observed metabolites from prokaryotic, plant, and fungal origin (the latter two were discarded during the curation). Many of the metabolites from non-diatomic sources were also marine natural products suggesting that both fungi and diatoms may produce similar compounds.

However, the annotation of certain spectral features to compounds of fungal origin could also be due to misannotation attributed to the limitations of mass spectrometry techniques and current annotation tools and databases, and the complexity of chemical data. Mass spectrometry (MS) currently is the main viable option for untargeted metabolomics followed by manual curation and verification [66]. The MS-based search for metabolite annotation rarely leads to unique biochemical structures due to the presence of compound isomers in the databases used for dereplication and the limitations in the mass accuracy of the MS system used [67]. The MS^1^ peaks chosen for MS^2^ fragmentation have varying optimal fragmentation energy, so not all MS^2^ spectra are informative. Features obtained with an untargeted LC-MS^2^ approach, which do not have an MSI level 1 confidence score (based on the known internal standard measurement), cannot provide absolute certainty about the assigned chemical structure for a feature due to the lack of our current knowledge of the chemical dark space. Moreover the current databases also only favor well-studied model organisms and may not cover the entire metabolome of *S. marinoi*, which is why it is important to elucidate the metabolome of this diatom.

To overcome a fraction of these challenges, MAW automatised the annotation process and performed dereplication in both spectral and compound databases while integrating prior biochemical knowledge in the form of a suspect list, which is also important to make accurate annotations. To give each annotation a confidence score, the Metabolomics Standards Initiative (MSI) levels of confidence were implemented in MAW to follow the standards set by the metabolomics scientific community. Automated workflows such as MAW, aid in the standardisation, and reproducibility and reduce the manual work in the analysis. Metabolomics is a relatively new omics field and the standardisation in metabolomics is not widely adopted which leads to missing metadata and reproducibility issues. However, it’s challenging to follow a set of standards in metabolomics because of the different combinations of chromatography and data acquisition methods of a diverse set of metabolites. Moreover, although using ChemONT, which is a standardised vocabulary for chemical classification, one metabolite with a carboxylic group and an amine group can be categorised as either an organic acid or an organic nitrogen compound. As of now, MAW follows the currently available standards, and ontologies for the spectral data acquired from data-dependent acquisition (DDA) LC-MS^2^ data, associating one chemical class to each annotation.

### 3.4 Future perspective

The catalogue of metabolites presented in this study exhibits a wide range of coverage provided by untargeted LC-MS-metabolomics. However, this approach does not provide validity for structure identification. To overcome this limitation, it is recommended to conduct targeted studies using some of the biologically important NPs. To further improve the accuracy of metabolite annotation in future studies, techniques such as nuclear magnetic resonance (NMR) spectroscopy can be employed to provide additional structural information for metabolite identification. The development of more comprehensive annotation tools and submission of new experimental spectra to natural product databases like GNPS will facilitate better metabolite annotation and identification, contributing to the expansion of available metabolites. Re-annotation with new updates of the databases can also enhance the annotation results. An integrative study of genomics and metabolomics from this diatom can also reveal the production of secondary metabolites involved in different biomechanisms in the marine ecosystem providing a systems-level understanding. Being the model organism for diatoms, this could hold the potential for elucidating biochemical mechanisms in other diatom species and unveiling marine-centric metabolic pathways [68].

## 4 Methods and Materials

### 4.1 Suspect List Development and Curation

For this study, a suspect list was generated manually from different databases and literary resources. The list contains metabolites that have been reported in *S. marinoi* and *S. costatum*. The 893 compounds in the list were collected and assembled from the databases such as KEGG [69–71], ChEBI [72], PubChem [73], MetaCyc [74], UniProtKB [75], BRENDA [76], LOTUS [77], and several publications [12, 16, 18, 35, 78–93]. For curation and additional metadata on each entry, PubChemPy [94], and RDKit [95], were used by adding any missing synonyms, IUPAC names, and molecular weights. Entries with no identifier or structural notation were discarded. Lastly, pybatchclassyfire [96] was used to add classification based on ChemONT 2.1 (subclass, class, superclass) to the suspect list using SMILES [23].

### 4.2 Sample preparation and metabolite extraction

For the whole metabolome extraction from LC-MS samples, methanol, ethanol, chloroform, acetonitrile, and water were used as reagents. S. marinoi RCC 75 strain used in this study was obtained from Roscoff Culture Collection (Roscoff, France). These cultures were grown in 40mL artificial seawater (ASW) inside 50 mL cell culture flasks. The cultures were kept at a standard temperature of 13 °C with 55-65 μmol photons m-2 s-1 lighting, keeping the 14:10 hr light:dark regime. The cultures were incubated in a shaking incubator at 80 rpm. Cultures were collected on Whatman GF/C filters with 1.2 μm pore size (GE Healthcare, US) under vacuum (750 mbar). These conditions provide a filtering speed of ∼1.5mL/min. The filters were then submerged in an ice-cold extraction mixture of methanol:ethanol:chloroform, 1:3:1, v:v:v. Samples were ultrasonicated for 10 min, and solutions were transferred to new Eppendorf tubes without the filters. The Eppendorf tubes with the solutions were then centrifuged at 30000g at 4 °C for 15 min. The aliquots for LC-MS analyses were separated and evaporated under a vacuum. Samples with aliquots that contained 3x106 cells were prepared. Media Blanks included the average volume for the culture’s samples. The volumes varied from 25 -40 μL depending on cell number in the cultures. The dried samples were stored in argon at -24 °C. For LC-MS analysis, the dried samples were dissolved in 150 μL of the mixture of methanol:acetonitrile:water, 5:9:1, v:v:v.

### 4.3 LC-MS^2^ Measurement

Samples separation with chromatography was performed on the SeQuant ZIC-HILIC column (5 μm, 200 Å, 150x2.1 mm, Merck, Germany) with a flow rate of 0.6 mL/min. The samples were measured with the Dionex Ultimate 3000 system coupled to a Q-Exactive Plus Orbitrap mass spectrometer (Thermo Scientific, Bremen, Germany). Using Electrospray ionisation, the molecular ions were obtained in both positive and negative modes, with a scan range of 80-1200 m/z. Samples were measured with a resolution of 70,000. The MS^2^ measurements were also performed on both modes, using parallel reaction monitoring (PRM), with a three-stepped normalised collision energy of 15, 30, and 45; a scan range of 80 -1200 m/z, and a resolution of 70,000. MS^1^ and MS^2^ measurements were performed in parallel. MS^2^ measurements were conducted with PRM using a target list of all detected compounds created with Compound Discoverer version 3.1 (Thermo Fisher Scientific). All MS^2^ spectra were acquired within 20 consecutive injections of the same sample.

### 4.4 Metabolic profiling with Metabolome Annotation Workflow (MAW)

The RAW MS files obtained from the Orbitrap Mass Spectrometer, were preprocessed using Compound Discoverer. The standard workflow (Untargeted Metabolomics with Statistics Detect Unknowns with ID Using Online Databases and mzLogic) was used with default options. The RAW MS^2^ files were converted to .mzML using Proteowizard MS-Convert Suite [97]. MAW (version 1.1) was used for dereplication against spectral and compound databases. For the spectral database dereplication, the input query spectra were matched against the experimental spectra from GNPS, HMDB, and MassBank. For the compound database dereplication, we used SIRIUS4 (version 4.9.12). SIRIUS performed the dereplication against the database “ALL” which refers to all databases integrated within the software. CANOPUS [34] and ClassyFire predicted the chemical class of the candidates. The candidate selection module was used to list the most probable (high-scoring) candidates from spectral databases and SIRIUS. The sunburst function generated sunburst diagrams based on the chemical classes of top-ranking candidates. Figure 3 explains the untargeted approach to extracting the whole metabolome of the diatom.

**Figure 3:**
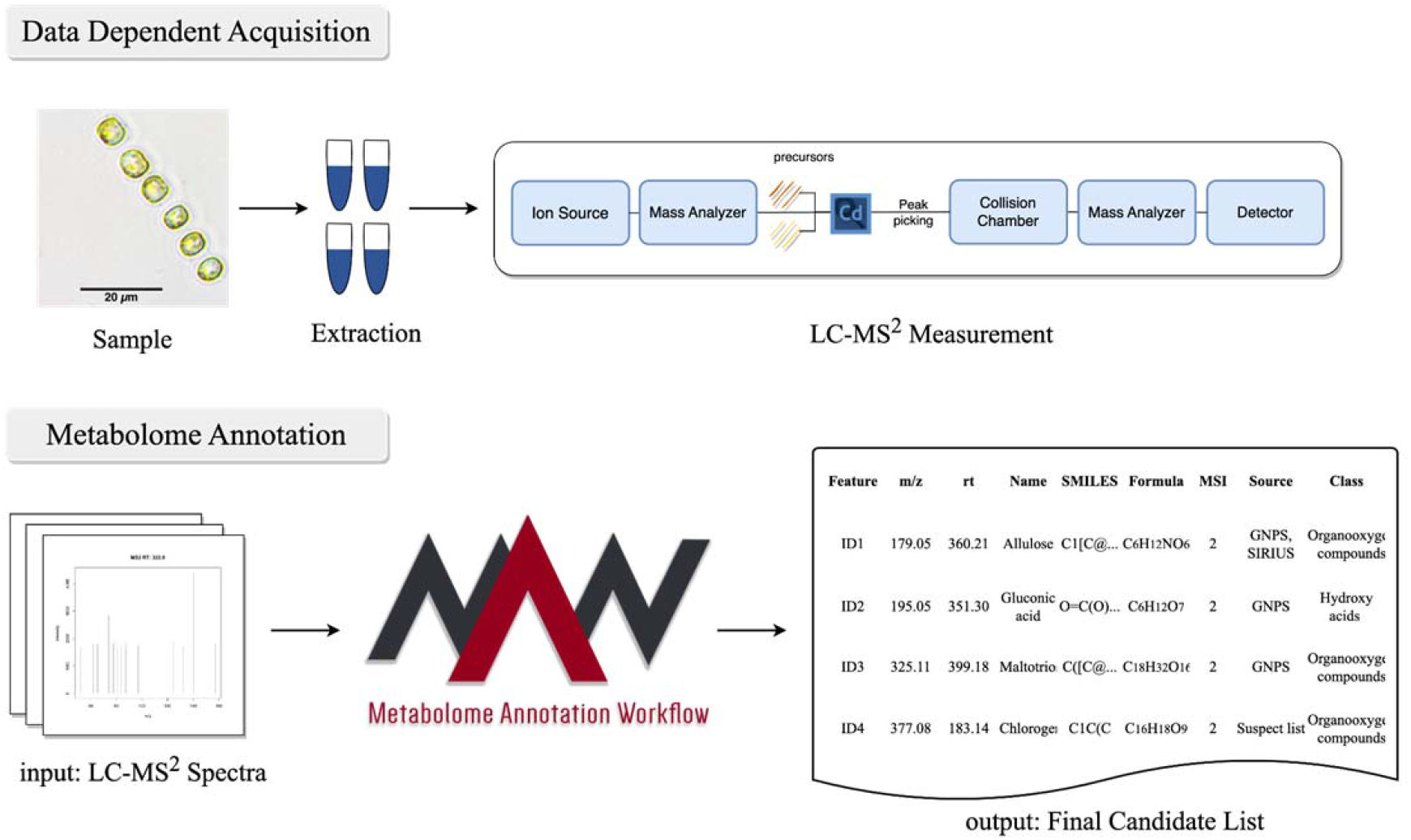
Workflow for untargeted mass spectrometry approach to acquire the whole metabolome from *Skeletonema marinoi*. After the sample collection and metabolic extraction, LC-MS^2^ measurements are performed using an orbitrap mass spectrometer. The data obtained from the LC-MS^2^ is then subjected to MAW which performs the annotation of metabolites and generates a final list of metabolites extracted from the samples.

### 4.5 Data Availability

The RAW and mzML metabolome data and the identifications are available on the MetaboLights repository [98] under study number MTBLS2892 (www.ebi.ac.uk/metabolights/MTBLS2892), the mzML files [References] are also available on Zenodo with the DOI: (pos) 10.5281/zenodo.7515842 (https://zenodo.org/record/7515842) [99] and (neg) 10.5281/zenodo.7515829 (https://zenodo.org/record/7515829) [100]. The currently available versions of the GNPS, HMDB, and MassBank are present on Zenodo with the DOI: 10.5281/zenodo.6528931 (https://zenodo.org/record/6528931) [101]. The suspect list for *Skeletonema* spp. is present on Zenodo with the DOI: 10.5281/zenodo.5772755 (https://zenodo.org/record/5772755) [102]. Scripts used to generate the list of metabolites using MAW version 1.1 and for the suspect list curation are available at https://github.com/zmahnoor14/MAW-Diatom [103]. The list of annotated metabolites from MAW, intersection, and union lists are available on Zenodo with the DOI: 10.5281/zenodo.7798782 (https://zenodo.org/record/7798782), along with the corresponding sunburst plots [104].

## Conflict of Interest

*The authors declare that the research was conducted in the absence of any commercial or financial relationships that could be construed as a potential conflict of interest*.

## Funding

Funded by the Deutsche Forschungsgemeinschaft (DFG, German Research Foundation) under Germany’s Excellence Strategy - EXC 2051 - Project-ID 390713860

## CRediT Author Statement

Mahnoor Zulfiqar: workflow execution and result analysis; writing the manuscript, worked on data availability, and curating the suspect list.

Daniel Stettin: performing LC-MS^1^ Analysis and LC-MS^2^ Measurements; writing methodology section.

Saskia Schmidt: generating the suspect list for *Skeletonema* spp. Vera Nikitashina: sample preparation and LC-MS^1^ measurements.

Georg Pohnert: provision of MS data, supervision of the project, obtaining the funds, and revising the draft.

Christoph Steinbeck: supervising the project, obtaining the funds, and revising the draft. Kristian Peters: supervising the project and revising the draft.

Maria Sorokina: Conception of the work, supervision of the project, and revising the draft. All authors reviewed the manuscript.

